# Blocking of EGFR Signaling is a Latent Strategy for the Improvement of Prognosis of HPV Induced Cancer

**DOI:** 10.1101/2020.10.06.329110

**Authors:** Jianfa Qiu, Feifei Hu, Tingting Shao, Yuqiang Guo, Zongmao Dai, Huanhuan Nie, Oluwatayo Israel Olasunkanmi, Yang Chen, Lexun Lin, Wenran Zhao, Zhaohua Zhong, Yan Wang

**Author notes:** Address correspondence to Yan Wang, and Zhaohua Zhong,. Jianfa Qiu and Feifei Hu contributed equally to this work. Author order was determined on the basis of their contribution.

## Abstract

Human papillomavirus (HPV) is a dsDNA virus and its high-risk subtypes increase cancer risks. Yet, the mechanism of HPV infection and pathogenesis still remain unclear. Therefore, understanding the molecular mechanisms, and the pathogenesis of HPV are crucial in the prevention of HPV related cancers. In this study, we analyzed cervix squamous cell carcinoma (CESC) and head and neck carcinoma (HNSC) combined data to investigate various HPV induced cancer common feature. We showed that epidermal growth factor receptor (EGFR) was downregulated in HPV positive (HPV+) cancer, and that HPV+ cancer patients exhibited better prognosis than HPV negative (HPV−) cancer patients. Our study also showed that TP53 mutation rate is lower in HPV+ cancer than in HPV− cancer and that TP53 can be modulated by HPV E7 protein. However, there was no significant difference in the expression of wildtype TP53 in both groups. Subsequently, we constructed HPV-human interaction network and found that EGFR is a critical factor. From the network, we also noticed that EGFR is regulated by HPV E7 protein and hsa-miR-944. Moreover, while phosphorylated EGFR is associated with a worse prognosis, EGFR total express level is not significantly correlated with prognosis. This indicates that EGFR activation will induce a worse outcome in HPV+ cancer patients. Further enrichment analysis showed that EGFR downstream pathway and cancer relative pathway are diversely activated in HPV+ cancer and HPV− cancer. In summary, HPV E7 protein downregulates EGFR that downregulates phosphorylated EGFR and inhibit EGFR related pathways which in turn and consequently induce better prognosis.

**Importance:** Although HPV infection has been studied in various cancer types, there are only limited studies that have focused on the common effect of HPV related cancer. Consequently, this study focused on CESC and HNSC, two cancer types with high HPV infection proportion in cohort, thereby, intending to dig out the common effects and mechanisms of HPV+ cancers.

Unlike some virus-human interaction prediction studies, the P-HIPSter database provides virus-human protein interaction based on protein structure prediction. Through this data, our interaction network was able to uncover previously unnoticed protein interactions. Our finding revealed that HPV infection caused various gene expression differences, and a great amount of which interact with EGFR, a cancer related gene. Therefore, since EGFR is associated with HPV+ cancer patients’ survival, some FDA proved EGFR inhibitors would be potential anti-HPV drugs.

## Introduction

Tumor can be caused by several factors (1). A virus is a small pathogen that often cause pathological changes or diseases in the target host (2). Some viral infections have been linked to be essential factors that induce numerous forms of cancer such as liver cancer and nasopharyngeal carcinoma (3, 4). Virus lifecycle requires intracellular environment owing to its simple structure (5, 6). It hijacks the cell’s complex protein and nucleic acid synthesis system for self-proliferation and also controls cells’ functional protein to modulate the normal cell signaling pathway (7).

Human papillomavirus (HPV) is a non-enveloped dsDNA tumor virus. Almost all cervical squamous cell carcinoma and about 40% of head and neck cancers are consequences of HPV infection (8, 9). HPV preferably infect the mucosal layer and no evidence shows that HPV has the ability to infect other cells except basal cells of the epithelia. Basal cell is the high differential ability cell of the epithelia. Hence, host cell development and differentiation ability are probably required for HPV infection (10). The carcinogenesis of squamous cell carcinoma is often accompanied by changes in development-related functions (11). Therefore, epithelia development regulated protein may be the key target of HPV infection and oncogenesis. HPV genome encodes seven early phase proteins (E1 to E7) and two late phase proteins (L1 and L2) for its proliferation. E6 and E7 proteins can modulate p53 and Rb through downregulation or inhibition, which is the basic mechanism of HPV+ cancer genesis (9). Therefore, E6 and E7 can be regarded as the most essential HPV oncogenic proteins (12). Due to its small genome and limited virus encoded protein, virus proteins require high efficiency and multifunctionality for complicated manipulation. For example, evidences showed that E6 and E7 proteins can interact with many human proteins and participate in a lot of biological processes (13). Likewise, HPV capsid protein L1 and L2 have been reported to interact with human proteins (14).

Many studies have examined HPV infection features in several types of cancers and HPV infection induced cancers have also been sufficiently investigated. For example, over 90% of the occurrence of CESC is attributed to HPV infection (15). Also, HNSC is highly linked to HPV infection (16). EGFR is a cancer-related gene and it also has been reported to be a potential biomarker of HPV infection. There are also studies that have demonstrated that some subtypes of HNSC exhibit higher HPV infection rate than other subtypes, and that HPV copy number is lower in HPV infected subtypes (17). Furthermore, EGFR is associated with HPV-related cancer prognosis. It was reported that EGFR and pEGFR are potential biomarkers of prognosis for oropharyngeal cancer (18). In cervical cancer, EGFR signaling can be affected by Hippo/YAP pathway and eventually influence cancer progression (19). Some reports suggested that EGFR expression can be regulated by HPV E5 protein (20), while contradicting reports showed that E5 protein does not regulate the expression of EGFR and cancer prognosis (21). Other reports showed that EGFR can be possibly regulated by miRNA. For example, in HPV-infected HuR cell, smoking induced control of miR-133a-3p regulates the expression of EGFR (22). Hence, EGFR expression in HPV-infected cancer may be regulated by multiple factors such as existing complex mechanisms and HPV viral protein. However, no study has completely established the HPV protein is the key modulator of EGFR.

Since most of the previous studies focused on comparing single cancer type or HPV+ groups with normal group. There are limited studies that focused on multiple cancer types or compared HPV+ cancer against HPV− cancer. Therefore, it is noteworthy to investigate EGFR regulating mechanisms in a multiple cancer types. Owing to existing EGFR targeted drugs, EGFR would be a potential target for HPV induced cancer prognosis improvement.

Our study analyzed CESC and HNSC combined data at multiple levels including mRNA, miRNA, SNV, and protein expression level. We also constructed a global network with HPV proteins, HPV differentially expressed genes and miRNAs in HPV+ cancers versus HPV− cancers. Through the network, our study showed that EGFR is regulated by HPV E7 protein and downregulated by miR-944. Furthermore, our findings showed that pEGFR and its up- and downstream proteins activation are negatively correlated with HPV+ cancer survival. These findings are evidences that EGFR is regulated in a complex mechanism and that E7 is the HPV protein that regulates EGFR expression in HPV induced cancer.

## Results

### HPV positive cancer patients are significantly different from HPV negative cancer patients in gene expression and they show higher survival possibility

In order to understand the relationships between HPV regulation and cancer, we selected the two most common HPV-related cancers which are CESC and HNSC for combined analysis. For RNA-Seq read counts matrix from ICGC database, principal component analysis (PCA) showed sample distribution of HPV+ samples and HPV− samples (Fig 1A). After removing the outlier at the lower right area, PCA plot was redrawn as displayed in Fig 1B, in which, HPV+ samples showed different distribution patterns against HPV− samples. The different distribution pattern shows that HPV+ cancer is distinct from HPV− cancer in gene expression. Further differentially expressed gene analysis was carried out by grouping samples by their HPV infection status. 834 differentially expressed genes were screened and these genes basically distinguished HPV+ from HPV− samples (Fig 1C). Survival analysis based on clinical data from FireBrowse database showed that HPV+ patients had better prognosis compared with HPV− patients (Fig 1D). It implies that different survival rate is attributed to DEGs to some degree.

**Fig 1.**
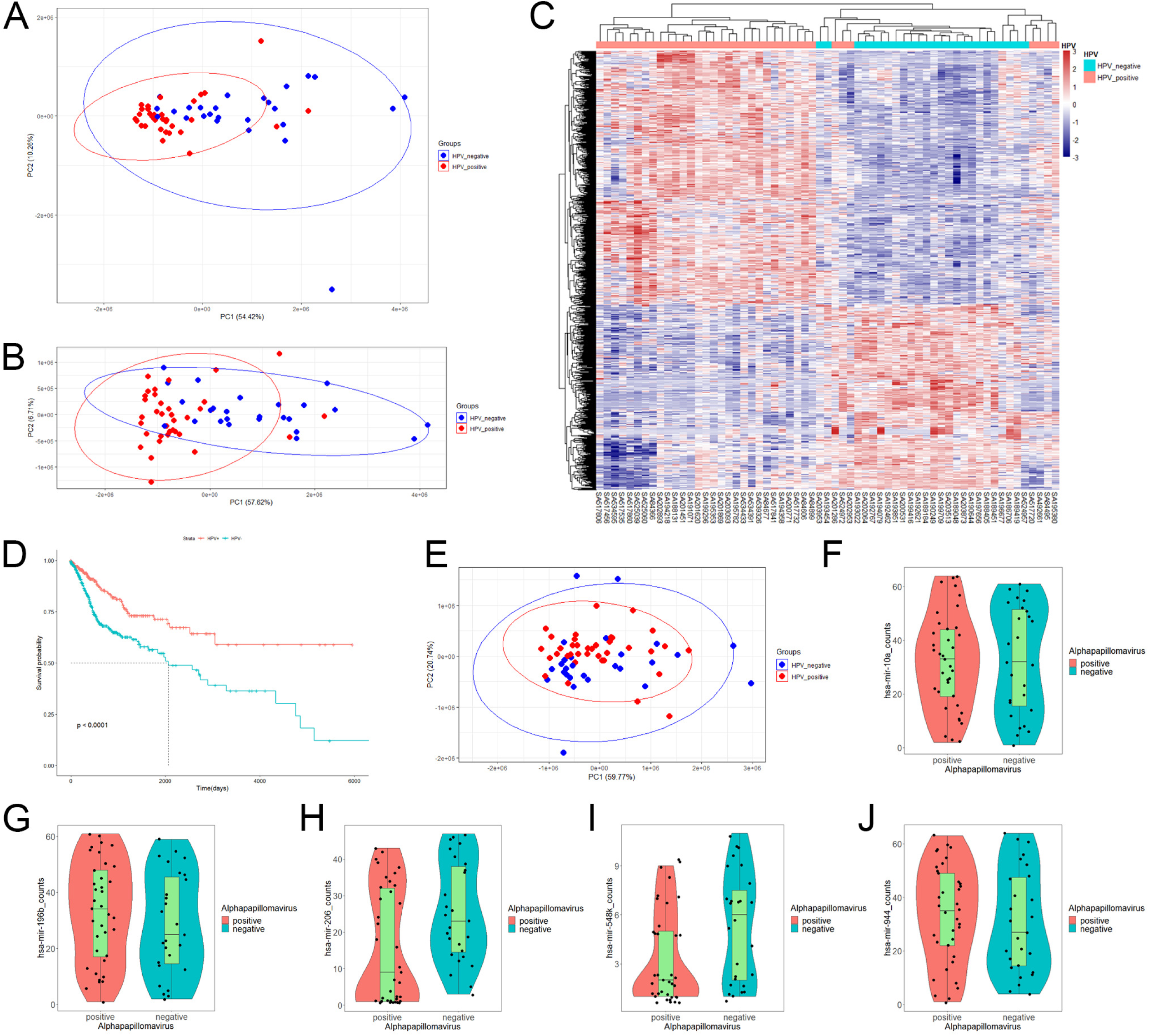
mRNA expression, miRNA expression and prognosis differences in HPV+ cancers against HPV− cancers. (A) Principal component analysis (PCA) plot of CESC and HNSC samples’ RNA-Seq data within PCAWG program of ICGC database. HPV+ samples are marked in red, HPV− samples are marked in blue. (B) Redrawn PCA plot after outlier was removed. HPV+ samples are marked in red, HPV− samples are marked in blue. (C) Differentially expressed genes (DEGs) heatmap of HPV+ group vs HPV− group. Gene FPKMs were scaled with z-score by samples. Pearson correlation coefficients were taken for samples and genes clustering. (D) KM-plot of HPV+ patients and HPV− patient survival status from TCGA CESC and HNSC project. (E) PCA plot of CESC and HNSC samples’ miRNA-Seq data within PCAWG program of ICGC database. HPV+ samples are marked in red, HPV− samples are marked in blue. (F-J) hsa-miR-10a, hsa-miR-196b, hsa-miR-206, hsa-miR-548k and hsa-miR-944 expression status in HPV+ samples and HPV− samples respectively.

The miRNA-Seq read counts data were also analyzed using PCA and differential expression analysis. PCA distribution showed no obvious difference between HPV+ and HPV− samples (Fig 1E), which indicates that there is no clear difference in miRNA expression level between HPV+ cancer and HPV− cancer. Differentially express analysis further showed that only 5 miRNAs were significantly differentially expressed. They were hsa-miR-944, hsa-miR-196, hsa-miR-206, hsa-miR-10a and hsa-miR-548k (Fig 1F–J). Further screening of differentially expressed miRNAs’ target in intersection, showed only hsa-miR-944, hsa-miR-206 and hsa-miR-548k DEG targets, which signifies that hsa-miR-196 and hsa-miR-10a probably does not participate in DEGs related functions although they were differentially expressed. Both hsa-miR-196 and hsa-miR-10a were upregulated in HPV+ cancers. The differential expression may be related to HPV proliferation. Small miRNA express difference between HPV+ cancer and HPV− cancer shows that only few miRNAs participate in HPV infection specific regulation and most of them are only cancer related or are steadily expressed in both situations.

### TP53 mutation proportion is lower in HPV+ cancer than in HPV− cancer

Single nucleotide variation (SNV) analysis was done for CESC and HNSC combined data. Comparing HPV+ and HPV− groups, the number of samples was almost the same. (Fig 2A). In determining the nucleotide variation type, we showed that HPV+ samples variation types and rates differs from that of HPV− samples. It showed that HPV+ samples’ C>G mutation rates are higher when compared with C>A mutation, while they are almost the same in HPV− samples (Fig 2B and 2C). At gene level for all samples, TP53 ranked at the 2nd place for single nucleotide mutation for all the genes. Moreover, TP53 mutation takes up 37% samples of the total mutation and it got the first place of all matched genes (Fig 2D). These results suggest that TP53 mutation plays a critical role in CESC and HNSC.

**Fig 2.**
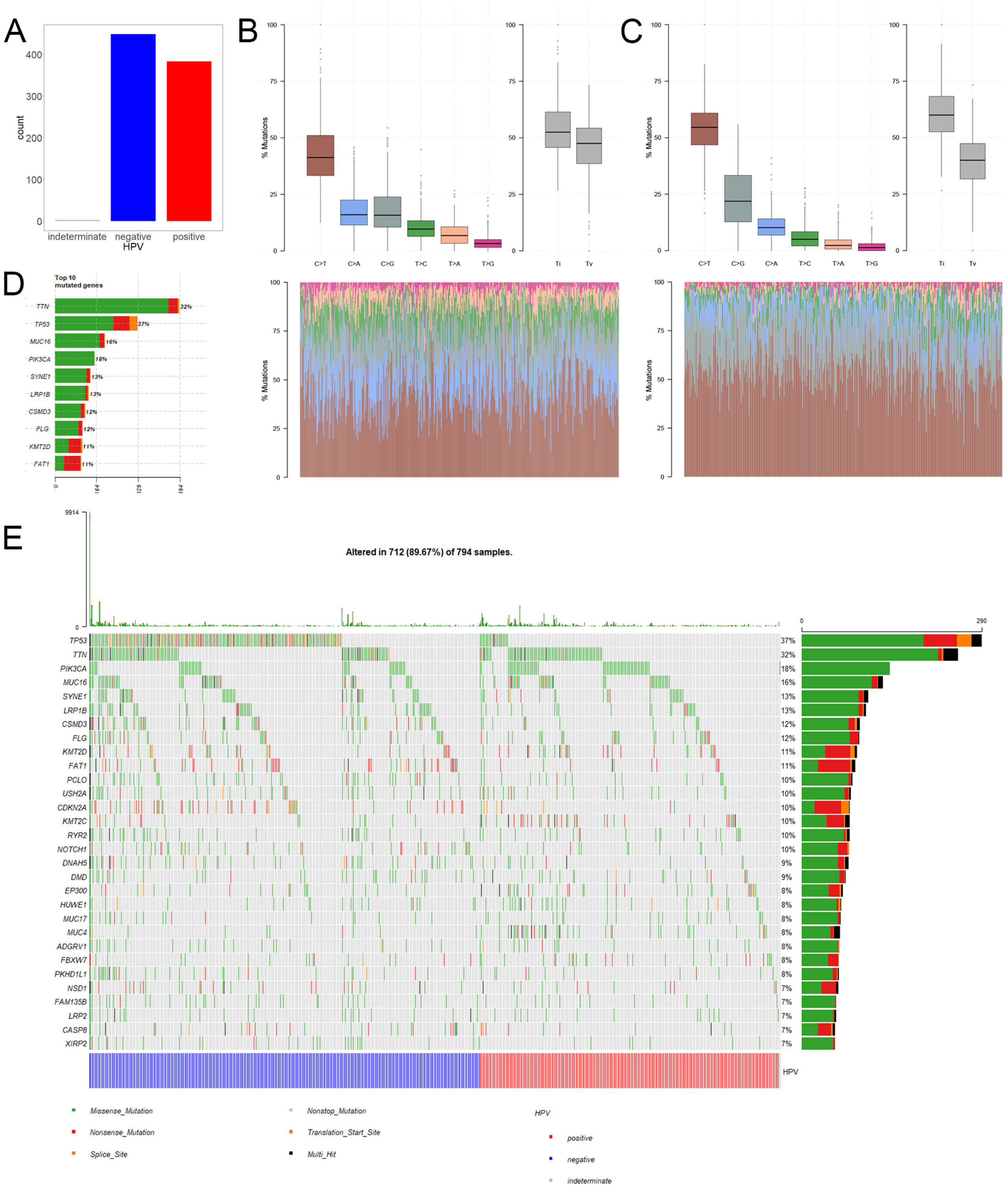
Single nucleotide variation (SNV) in HPV+ cancers and HPV− cancers. (A) Total numbers of HPV+ samples and HPV− samples in SNV data of TCGA CESC and HNSC project. MUSE software processed SNV data was used in our study. (B) HPV+ samples single nucleotide mutation type proportions. (C) HPV− samples single nucleotide mutation type proportions. (D) Top 10 highly mutated genes’ mutation rates and mutated sample counts. (E) Oncoplot of the top 30 highly mutated genes, samples were grouped by HPV infection status.

Furthermore, we selected top 30 genes with the highest mutation frequency and used oncoplot to show their mutation rates in each of the samples. The result showed that the number of TP53 mutation is significantly higher in HPV− cancer than in HPV+ cancer. Whereas most of the genes with high mutation rate showed no significant difference in both groups (Fig 2E). This result suggests that TP53 mutation is an important mechanism for the occurrence of HPV−. However, HPV+ cancer shows no relative involvement with TP53 mutation, cancer occurrence may be involved in other mechanisms.

### HPV protein regulation of human protein is an important mechanism for the occurrence of HPV+ cancer

With virus protein-human protein, mRNA-mRNA (differentially expressed) and miRNA-mRNA interaction pairs, we constructed a miRNA-mRNA-protein interaction network. The result showed that a considerable number of human genes are regulated by HPV proteins. Thus, the genes that are modulated by HPV viral protein may possibly be the key factors that induce tumor occurrence. (Fig 3A). Overall, most of the genes directly regulated by HPV are not differentially expressed gene. This suggests that HPV-regulated genes have similar expression pattern in HPV− tumors.

**Fig 3.**
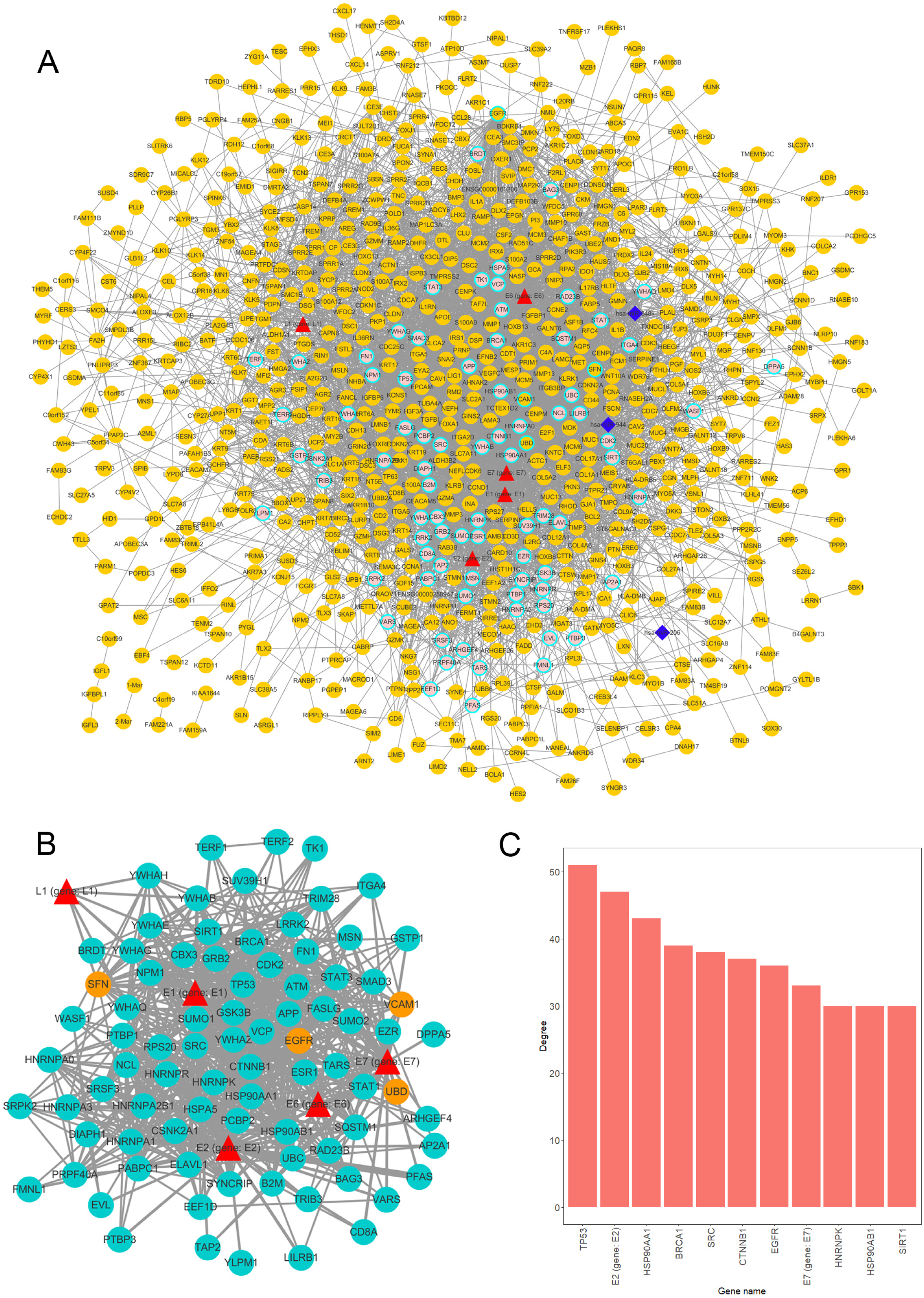
Overview of interaction network. (A) Global network of miRNA-mRNA-protein interactions. HPV proteins are marked as red triangles; miRNAs are represented as deep blue diamonds; differentially expressed genes (DEGs) are orange circle; non-differentially expressed genes are in pink; Circles with light blue border are HPV directly interacted human genes. (B) HPV proteins directly regulated subnetwork. Turquoise circles are HPV directly manipulated non-differentially expressed genes; red triangles are HPV proteins; orange circles are HPV directly manipulated DEGs. Top 11 genes degree distributions of HPV proteins directly regulated subnetwork.

We further extracted a subnetwork that contains only HPV protein and its regulated genes to investigate the manipulation details. From the subnetwork, 4 genes were differentially expressed and are listed as follows, EGFR, SNF, UBD and VCAM1 (Fig 3B). Further findings indicated that the variation between HPV+ cancer and HPV− cancer can possibly be attributed to the effect of HPV regulation of those 4 genes. Through degree analysis of subnetwork, we showed that degrees of tumor suppressor gene TP53 (regulated by HPV E7 protein) is the highest of all genes (Fig 3C). This result indicates that the regulation of TP53 by HPV is a crucial mechanism for HPV+ tumor to occur.

### EGFR is the crucial gene that regulate HPV+ tumor differentially expressed genes

We used degree = 55 to screen hub nodes of miRNA-mRNA-protein network, and 22 nodes were selected. Out of the 22 genes, 8 genes are DEGs. While 15 genes are direct HPV regulated genes, and nodes like TP53, BRCA1, EGFR and CTNNB1 are classic tumor related genes (Fig 4A). Remarkably, EGFR is the only hub node that belongs to both DEGs and directly interacts with HPV (Fig 4B). We further extracted a subnetwork constructed with EGFR and its first neighbor. The result showed that EGFR interacts with a considerable amount of DEGs and it is also regulated by hsa-miR-944 and HPV protein E7 (Fig 4C). This indicates that EGFR is an essential gene that regulates the differentially expressed genes.

**Fig 4.**
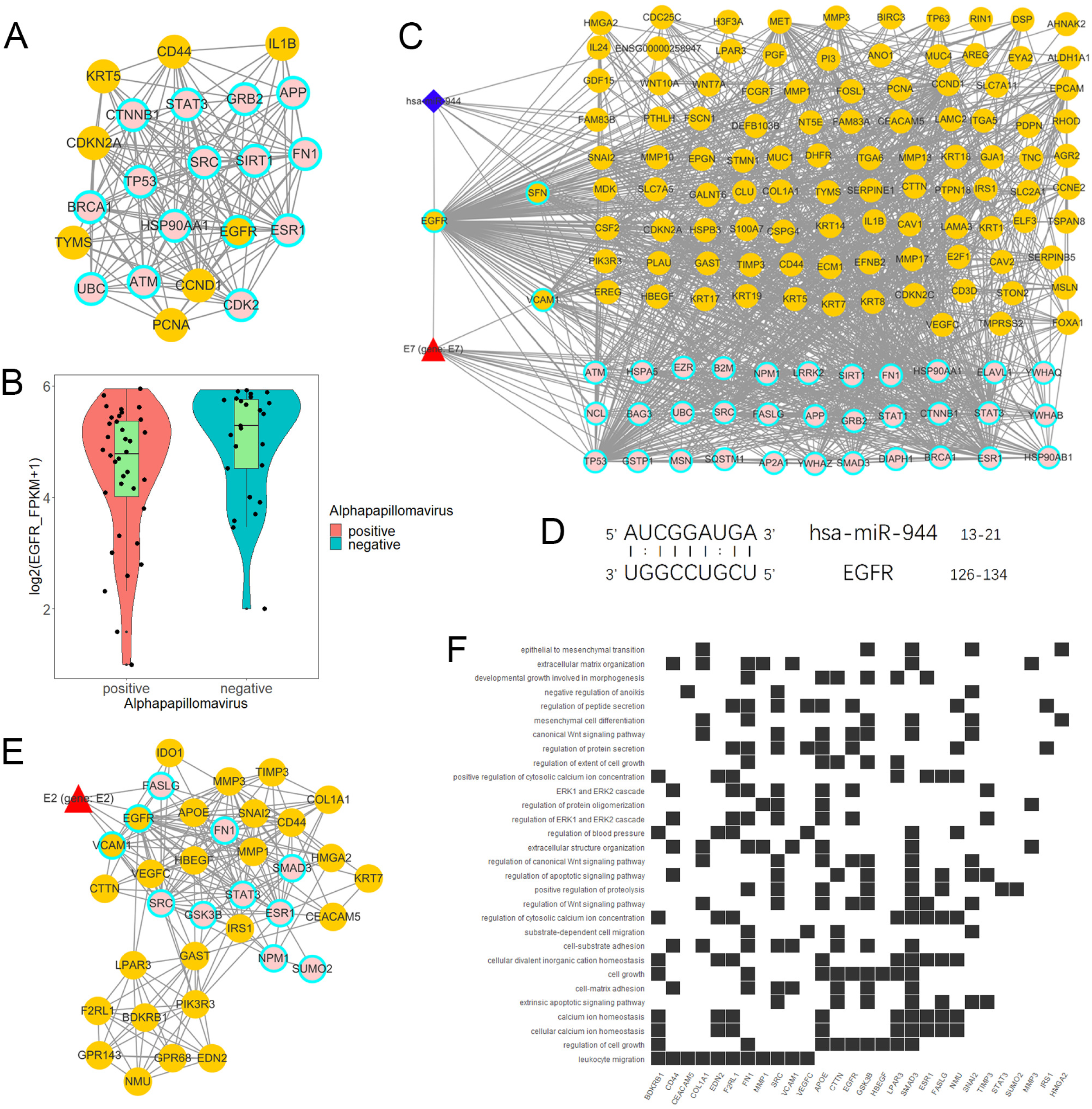
Analysis of EGFR related network. (A) Hub genes of global network, including 15 HPV manipulated genes; 7 DEGs are not regulated by HPV and 1 HPV regulated DEG. (B) EGFR express status in HPV+ samples and HPV-samples respectively according to RNA-Seq FPKM from ICGC PCAWG project CESC and HNSC data. (C) Subnetwork of EGFR first neighbors. (D) EGFR RNA binding prediction site with hsa-miR-944 using RIsearch2 software. (E) MCODE calculated EGFR containing network cluster. (F) EGFR containing cluster GO and KEGG enrichment analysis.

Furthermore, we predicted that EGFR is regulated by hsa-miR-944; and that the upregulation of has-miR-944 caused the downregulation of EGFR (Fig 4B and 1J). In order to confirm whether hsa-miR-944 combine stably with EGFR, RIsearch2 software was used for RNA combined analysis. The result shows that EGFR and hsa-miR-944 have low energy binding site at 3’ end of hsa-miR-944 (Fig 4D). Our findings revealed that EGFR is regulated by both E7 protein, and hsa-miR-944. This shows that E7 protein does not only induces carcinogenesis in HPV+ tissues, but also causes the difference in appearance in HPV+ and HPV− tumor.

A module with EGFR was identified using module analysis of global network. Since gene in the same module interact closely, there is possibility that they can participate in the same biological process. HPV protein E2, a key protein that plays a pivotal role in HPV infection from the early stage to the late stage, was also identified (Fig 4E). This finding suggests that EGFR participates in all HPV infection stages and could probably influence tumor development and prognosis. Likewise, GO and KEGG enrichment analysis revealed that the module genes are enriched with EGFR downstream pathways and participates in several functions, including development regulation, epithelial regulation and MAPK pathway (Fig 4F). This implies that EGFR downstream pathways are regulated by E2 and it can influence oncogenesis at late stage of infection.

### Activation of EGFR-related pathway is an important factor that decrease survival

In order to figure out how EGFR influence prognosis, we merged RPPA data of CESC and HNSC and a total of 133 protein was obtained. Student-t test was used to test for the differences between HPV+ and HPV− groups. Interestingly, 123 proteins were differentially expressed and only 10 proteins were not significant. Yet, TP53 was not differentially expressed at the protein level. Subsequently, COX proportional hazard regression model was used to evaluate the correlation between survival time and deferentially expressed proteins in HPV+ tumor patients. The results suggest that the expressed level of EGFR did not have any significant relationship with prognosis. Notably, there was a negative correlation between all tyrosine residue phosphorylated forms of EGFR and patients’ survival. Phosphorylation on site pY1068 and pY1173 representing EGFR was activated to form a dimer that binds with its ligand; thus, further activating downstream pathways like PI3K/Akt, MAPK, WNT pathway. We also highlighted that AR (amphiregulin), an EGFR ligand, and some significant proteins belongs to PI3K/Akt or MAPK pathway. Those downstream proteins also showed similar properties of phosphorylated form of EGFR that are significantly negatively correlated with survival (Fig 5).

**Fig 5.**
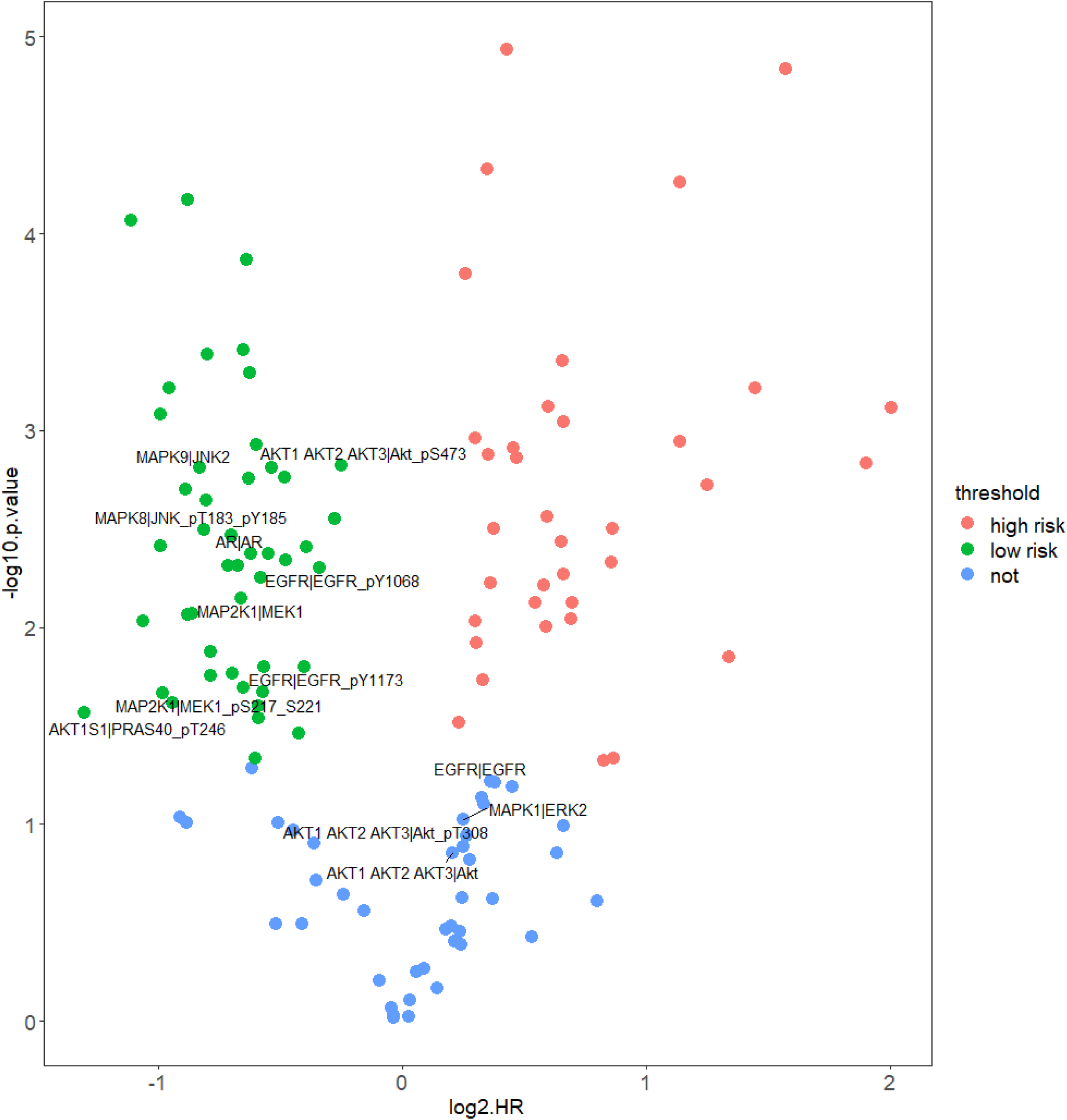
Correlation of protein expression and HPV+ patients’ survival. Figure shows COX proportional hazards regression. It measured correlation significance of protein expression and HPV+ patients’ survival. Only differentially expressed proteins (DEPs) of HPV+ vs HPV− are displayed. Proteins which *p* < 0.05 are significantly correlated with HPV+ patients’ survival, log2HR > 0 is positively correlated with survival, log2HR < 0 is negatively correlated with survival.

### EGFR modulates cancer prognosis through the regulation of immune and DNA repair pathway

In order to further find out the prognosis related pathways, we carried out KEGG enrichment analysis of HPV+ versus HPV− DEGs and mRNA targets of DEmiRs. The top enriched term list showed “human papillomavirus infection”, demonstrating that HPV activates a unique pathway different from HPV− cancer. For DEGs enrichments analysis, immune related pathway (such as IL-17 signaling pathway and TNF signaling pathway), cancer related pathway, and DNA related pathway were shown and it could be correlated with prognosis. Remarkably, EGFR related pathways, “PI3K/Akt pathway” and “ECM-receptor interaction”, were also included in the top list (Fig 6A). For DEmiR target enrichments, numerous of cancer related pathway were shown and EGFR related pathway, “PI3K/Akt pathway”, was also enriched (Fig 6B). These results suggest that HPV+ cancer show different prognosis related pathway activation compared to HPV− cancer, even in cancer related pathway. Gene Set Enrichment Analysis (GSEA) further showed that immune-related pathways, and tumor-related pathways were inactivated and DNA repair pathways were activated in HPV+ cancer groups compared HPV− groups. All these activated and inactivated pathways can possibly enhance better prognosis of HPV+ cancer (Fig 6C–K). Since EGFR is the hub gene of DEGs network, EGFR can possibly affect the prognosis of tumors through the regulation of immune, tumor and DNA repair pathway.

**Fig 6.**
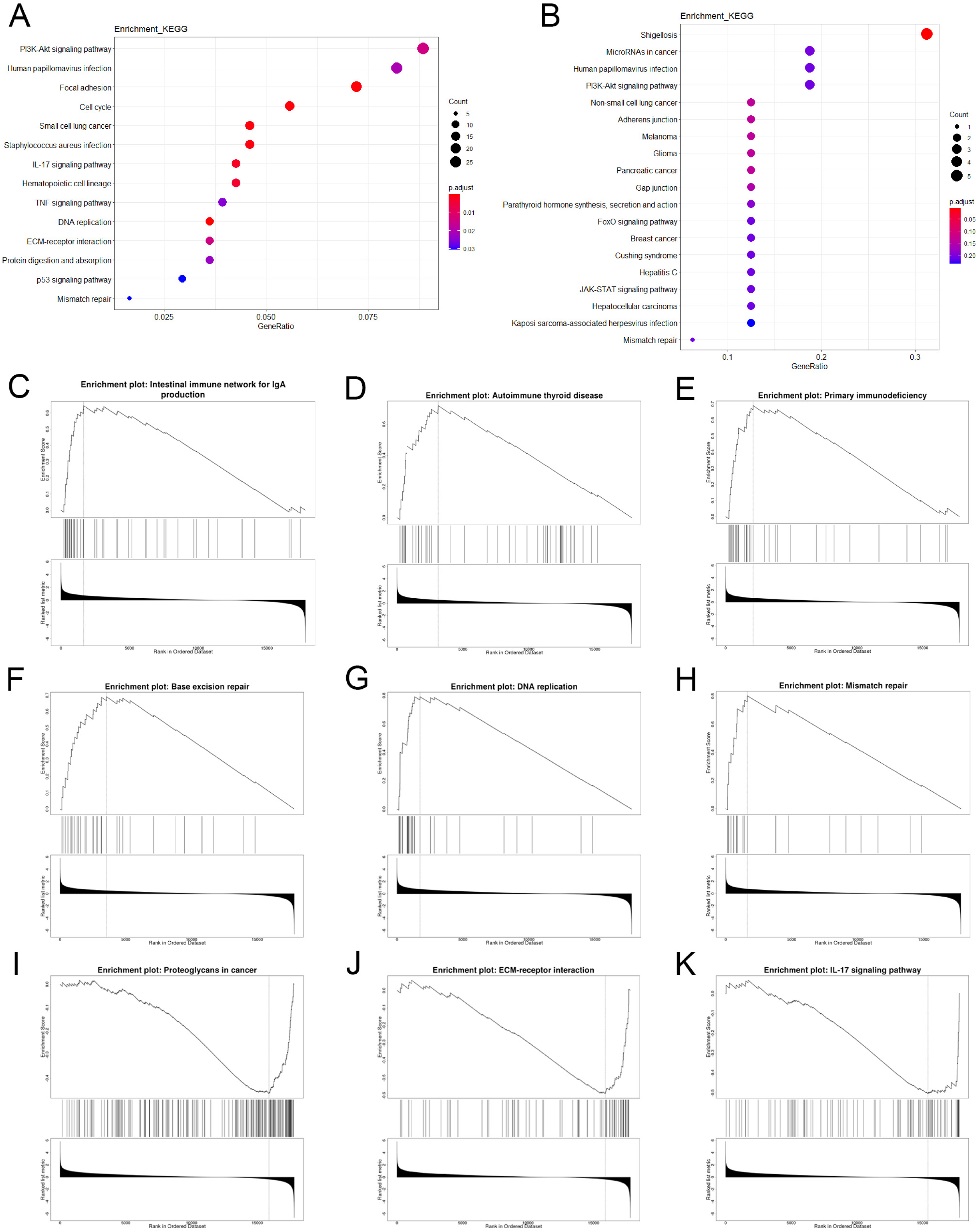
Enrichment analysis of mRNA and miRNA targets. (A) and (B) are top KEGG enriched terms of differentially expressed genes and differentially expressed miRNA target genes respectively. (C-K) are GSEA enriched terms, in which genes were sorted by log2FC.

## Discussion

HPV is a tissue specific oncogenic virus that specifically infect epithelium tissues. The mechanisms of HPV-induced squamous cell carcinoma are probably not similar to the mechanisms of HPV non-associated squamous cell carcinoma. Since both mechanisms cause tumorigenesis, it suggests that their gene expression patterns are somewhat common. Based on this hypothesis, our study compared HPV+ and HPV− tumors at multi-omics level in order to identify similar and different underlining molecular mechanisms. Although HPV can infect various type of epidermal tissues, CESC and HNSC are the most common type of HPV induced squamous cell carcinoma. Considering the representative of data and limited data volume, we combined both CESC and HNSC data into our study. Also, since our study focus on HPV-induced tumorigenesis rather than ontogenesis, we merged the two cancer data for analysis instead of separate analysis.

Recent reports revealed that HPV+ patients shows better prognosis than HPV− patients in certain types of tumor (23). And, our study is consistent with this result despite merging data from two different types of cancer. It implies that the better prognosis of HPV+ tumor is consistent across different tumor types. This effect is probably related to the difference in the expression pattern of prognosis associated genes. Although, different in mechanisms, both HPV− and HPV+ tumors express similar pattern in tumor associated genes. Virus infection influence host gene expression pattern mainly through direct regulation by virus-derived proteins and this process rarely induce gene mutations (24). The tumors that are not driven by virus usually shows mutations at oncogenes and/or tumor suppressor genes. In the total of 84 HPV-regulated genes, only 4 genes (EGFR, SNF, UBD and VCAM1) were differentially expressed when compared with HPV− tumor. Among the 4 genes, SNF and VCAM1 directly interact with EGFR through string estimation, this suggests that HPV-regulated DEG tends to interact, and that they participate in similar biological processes that affects patient’s survival.

Our RNA-Seq PCA showed that HPV+ tumors are distributed in different area against HPV− tumor at the first principal component. Although, the dispersion within group is not obvious, we believe that HPV infection status probably influence gene expression more than the primary tumor site. The 834 DEGs obtained from differentially express analysis further confirms that gene expression patterns of HPV+ tumors are different from HPV− tumors. Although, the clustering analysis showed that the clustering of samples was basically the same as that of HPV infection, a small number of HPV+ and HPV− clustered together. There are two possible reasons for this phenomenon. First, the samples’ HPV expression level may be very low, which may induce their gene expression patterns closer to that of HPV− tumors. Second, these samples were probably infected by low-risk HPV that rarely induce cancer, which implies that their oncogenesis ability is relatively lower. In spite of diversity in mRNA expression, miRNA expression between HPV+ and HPV− shows little differences. However, based on the enrichment analysis of DEmiRs target genes, we showed that there are several differences in tumor-related pathways of HPV+ and HPV− cancers. One possible reason for this is that only a small percentage of miRNAs are differentially expressed in HPV+ and HPV− tumors. And these differentially expressed miRNA are highly correlated with tumor-related biological processes.

Since certain mutations occur across different cancer types (25, 26), we therefore focus on differences in mutation types between HPV+ and HPV− tumor. Our study showed that TP53 mutation rate in HPV+ tumor is dramatically lower than in HPV− tumor. TP53 is an important tumor suppressor gene. The mutation of tumor suppressor gene is considered more serious than its dysfunction. HPV+ cancer patients showed better outcome than HPV− cancer patients and that can probably be attribute to low TP53 mutation rate (9, 27). We also showed that genes that belong to the same family or participate in the same pathway have higher mutation rate. For instance, mucin glycoprotein encoded genes MUC16, MUC17 and MUC4 ranked among the top 30 mutated genes. This suggests that mucin glycoprotein mutation is a signature of HNSC and CESC (28, 29). Moreover, MUC4 mutation rate in HPV+ samples are obviously higher compared with HPV− samples. We believe that mucin glycoprotein subtype and their mutation rates could be a latent biomarker for tumor classification.

Virus protein expression and regulation of biological function are usually diverse in HPV infection stage. For HPV, functional proteins like E1, E2, E5, E6 and E7 are highly expressed at early infection stage. Using these early stage proteins, HPV can hijack DNA and protein synthesis machinery of the cell for self-proliferation. At the final stage, HPV capsid proteins, L1 and L2, are expressed for virus assembly and escape from the cell (30). Reports suggests that, E2, an early stage protein, possibly participate in the late stage of HPV replication by activating DNA damage response (31–33). Our result shows that, EGFR is regulated by HPV oncoprotein E7 and that it takes part in E2-related regulation unit. This suggests that EGFR is involved in E2-regulated DNA damage response (34). Also, DNA related functions of HPV+ cancers are significantly activated compared to that of HPV− cancers. Beside HPV proteins, differentially expressed miRNAs also participate in EGFR regulation. Although differentially expressed miRNAs and their targets are rare, it is not accidental that differentially expressed miRNA targets EGFR (Hypergeometric test *p* < 0.001).

Since EGFR is a potent oncogene, EGFR dysregulation will cause several forms of cancer. High proportion of non-small cell lung carcinomas express EGFR and the EGFR mutant as its signature (35, 36). Likewise, EGFR has become a biomarker of HNSC (37). In our study, EGFR shows different express pattern for HPV+ and HPV− cancers. EGFR is a critical receptor that transduce epithelium growth and developmental signal into the cells. It plays an important role in epithelial stem cell division and differentiation. HPV does not only infect the basal cells of epithelium tissues, but also require epithelial development for its replication. Hence, regulation of basal cell differentiation is a crucial control strategy for HPV replication. On the other hand, HPV− cancers do not require epithelium differentiation for its replication. Therefore, EGFR downregulation is possible a potential strategy that targets HPV specific lifecycle. Tyrosine phosphorylated EGFR is the activated form of EGFR. Only the activated form of EGFR can serve as a receptor and receive extracellular signals. Downregulation of the overall expressed EGFR can possibly decrease the expression of phosphorylated EGFR, thus inhibiting the downstream signaling pathway of EGFR (38).

In summary, HPV+ cancer is significantly different from HPV− cancer in many aspects like DNA mutation, mRNA and protein expression. The initiation of cancer in HPV+ cells results from the regulation of biological processes related to host development by viral proteins. In contrasts, HPV− cancer is activated by several categories of risk factors and its highly related to TP53 mutation. Although distinct in mechanisms, both HPV+ and HPV− cancers are triggered by onco-related genes dysregulation. This study showed that EGFR is possibly the core molecule that affect immune and cancer related biological processes and it can eventually cause prognosis differences between HPV+ and HPV− cancers.

## Materials and Methods

### Data source

CESC and HNSC RNA-Seq read counts data of PCAWG project were obtained from ICGC database (https://icgc.org/). In the PCAWG project, information on HPV infection in each of the samples was from a study recently published by Marc Zapatka et al (39). CaPSID, P-DIP and SEPATH pipelines were used for HPV reads detection from PCAWG samples in Marc’s study, and samples with HPV reads that were detected in at least 2 pipelines were considered infected. The cancer genomic atlas (TCGA) data of simple nucleotide variation (SNV) are from Genomic Data Commons Data portal (GDC, https://portal.gdc.cancer.gov/). The source of reverse-phase protein array (RPPA) level 3 data and clinical data of TCGA samples were obtained from FireBrowse database (http://firebrowse.org/).

### Principal component analysis

Principal Component Analysis (PCA) was applied for data dimension reduction. Samples distribution confidence interval of each sample groups were displayed. The samples located far away from confidence interval ellipse were considered as outliers and were deleted in the subsequent exploration.

### Differentially express analysis

Limma package of R was used for RNA-Seq data and miRNA-Seq data differentially express analysis. Differentially Expressed Genes (DEGs) and differentially expressed miRNAs (DEmiRs) were identified by *p.adjust.* < 0.05 and |log2FC| >1.2. BH method was applied for *p* value adjustment. Student-t test was used to analyze differentially expressed RPPA data. Proteins with *p.value* < 0.05 were regarded as differentially expressed proteins (DEPs).

### Survival analysis

Clinical data from FireBrowse database was used for the prediction in survival differences of the two groups. To compare the data from the HPV+ tumor patients and HPV− tumor patients, Kaplan-Meier method was used for survival rate prediction and Kaplan-Meier plot (KM plot) was used to determine the survival curve. COX proportional hazards regression model was used to predict the correlation between differentially expressed protein (from RPRA data) and HPV+ patients’ survival time. *p*<0.05 was considered to significantly correlate with survival time, HR>1 was considered as positive correlation with survival time and HR<1 was considered to be negative correlation with survival time.

### miRNA target prediction

TargetScan (http://www.targetscan.org/vert_72/) and miRDB (http://mirdb.org/) database were used for differentially expressed miRNA target prediction. Targets that appeared in both databases were considered well-predicted. RIsearch2 software was used for further verification of specific interesting miRNA-mRNA interaction.

### SNV analysis

Single Nucleotide Variation (SNV) data of CESC and HNSC project were downloaded from TCGA GDC data portal. The processed MAF data used in this study was downloaded from MUSE software processed. Maftools R package was used for MAF file data mining. Maftools was likewise used for statistical analysis of gene mutation and data visualization.

### HPV-miRNA-mRNA network construction and analysis

HPV-human protein interaction prediction data were downloaded from P-HIPSTer (http://phipster.org/), a database that predicts virus-human protein interactions based on structural information. Differentially expressed mRNA interactions were predicted by String database (https://string-db.org/). MCODE plugin of Cytoscape (version 3.7.0) was used for network module identification. The parameters for MCODE were set as default. The degree calculation was determined by NetworkAnalyzer, and genes with degree higher than the threshold were defined as hub genes.

### Enrichment analysis

KEGG pathway enrichment analysis were determined using R package clusterProfiler. Hypergeometric test was used for terms significance testing. The *p.adjust.* < 0.2 was set as the threshold of significantly enriched terms. Webtools WEBGESTALT (http://www.webgestalt.org/) was used for GSEA enrichment analysis and results visualization (40).

## Acknowledgements

This work was supported by the National Natural Science Foundation of China grants to Y.W. (81772188) and Z.Z. (81571999 and 81871652); Natural Science Foundation of Heilongjiang Province grant to L.L. (LH2019H004). Heilongjiang Postdoctoral Scientific Research Developmental Fund to L.L. (LBH-Q19146).

We thank Miss. Yue Qi for the helpful discussions and Mr. Yuxiang Pan for his encouragement.

J.Q., F.H. and Y.W. conceived the study. J.Q., F.H. and Y.W. designed the experiment. J.Q. performed the experiment and computational analysis. Y.G., Z.D., H.N. and Y.C. collected the data. J.Q. wrote the manuscript. O.I.O., W.Z. and L.L. provided valuable suggestion to improve the manuscript. T.S. and Z.Z. provided the professional consulting support.

